# ACE2 Phosphorylation Modulates Angiogenesis via the Activator Protein-1

**DOI:** 10.1101/2025.10.17.683185

**Authors:** Yuyang Lei, Yuanming Xing, Yi Lei, Qiangyun Liu, Tong-You Wade Wei, Baoyu Li, Fangzhou He, Ziru Wu, Lu Wang, Chen Wang, Wenyang Hao, Jin Zhang, Ying Xiong, Juan Zhou, Xiaohong Wang, John Y-J. Shyy, Zu-Yi Yuan

## Abstract

**Background:** Angiogenesis plays a crucial role in organ development. However, aberrant blood vessel growth is involved in various diseases, including tumors and neovascular eye diseases. Angiotensin-converting enzyme 2 (ACE2) is a critical enzyme regulating the health of cardiovascular system, and its post-translational modifications (PTMs) are crucial to determine ACE2 expression level and activity. Here, we studied how the PTM of ACE2 in vascular endothelial cells (ECs) affect pathological retinal neovascularization and tumor angiogenesis.

**Methods:** The angiogenic capabilities of ECs were assessed by tube formation, sprouting assays, and 5-ethynyl-2-deoxyuridine (EdU) and filopodia staining. EC angiogenesis was examined by poteome profiler array and aortic ring assays in vitro and by the oxygen-induced retinopathy (OIR) and tumor angiogenesis models in vivo. High-throughput screening involving data from RNA-seq, ATAC-seq, and ChIP-seq were used to explore the epigenetic and transcriptional regulations of pro-angiogenic genes regualtged by ACE2 PTMs.

**Results:** ACE2 Ser-680 dephosphorylation in connection with Lys-788 ubiquitination increased EC angiogenic phenotype, which were manifested by aberrant vascularization in mouse OIR and tumor models. ACE2 Ser-680 dephosphorylation led to the activation of activator protein-1 (AP-1), which transactivated multiple genes involved in angiogenesis. AP- 1 inhibition mitigated such angiogenesis in vivo.

**Conclusion:** Our findings show a novel PTM mechanism of ACE2 involved in pathological angiogeneis. Specifically, ACE2 Ser-680 dephosphorylation facilitated AP-1 transactivation of the downstream pro-angiogenic genes in ECs.

## Introduction

Angiogenesis plays a crucial role in organ development, but aberrant blood vessel growth is involved in various diseases during adulthood, including tumors and neovascular eye diseases.^1^ Pathological retinal neovascularization, characterized by an aberrantly proliferating vascular tuft structure, is a defining feature of retinopathy of prematurity (ROP) and proliferative diabetic retinopathy.^2,3^ During tumor development, angiogenesis refers to the formation of new blood vessels within solid tumors to supply nutrients and oxygen, thereby promoting the spread of tumor cells.^4^ This pathological process involving vascular endothelial cells (ECs) is critical for tumor metastasis.^5^ Evidence from recent studies has led to the proposal that inflammation can drive ocular neovascularization.^6,7^ Inflammation also predisposes to the development of cancer and promotes tumor angiogenesis.^8^ Although tremendous effort has been attributed to therapeutic strategies targeting angiogenesis, there is still an unmet need to identify new molecules and pathways that regulate both physiological and pathological vessel growth.

Angiotensin-converting enzyme 2 (ACE2), a crucial member of the renin–angiotensin system (RAS), is an indispensable regulator in cells and tissues in the cardiovascular system. Functioning as the protective arm of RAS, ACE2 catalytically converts angiotensin I (Ang I) to Ang (1–9) and Ang II to Ang (1–7).^9^ In ROP, the deleterious arm of RAS, including renin, angiotensinogen, ACE, and Ang II type 1 receptor (AT1R), is upregulated, which suggests a link between RAS dysregulation and ROP progression.^10^ Regarding RAS in tumorigenesis, the activation of AT1R enhances cancer cell migration, proliferation, angiogenesis, and tumor metastasis, including melanoma, ovarian carcinoma and renal cell carcinoma.^11–13^ In contrast, ACE2, along with Ang (1–7), has an inhibitory effect on angiogenesis in breast cancer and renal carcinoma.^14,15^

Post-translational modifications (PTMs) of ACE2 play a crucial role in determining its levels, activity, and functions. Protein phosphorylation and ubiquitination are two important PTM mechanisms. We previously demonstrated that AMP-activated protein kinase (AMPK) phosphorylation of ACE2 at Ser-680 suppresses its Lys-788 ubiquitination by murine double minute 2 (MDM2), thereby reducing proteasome-mediated degradation.^16,17^ This AMPK phosphorylation of ACE2 Ser-680 in ECs enhances ACE2 protein stability, Ang (1–7) production, and endothelial nitric oxide synthase-derived NO bioavailability.^16,17^ ACE2 seems to act against ROP and tumor angiogenesis, but how PTM regulations of ACE2 in ECs affect these pathophysiologic processes remains unknown.

Here, we report novel findings that ACE2 Ser-680 phosphorylation is decreased in murine oxygen-induced retinopathy (OIR) and melanoma models. Conversely, exogenously elevated ACE2 Ser-680 phosphorylation could suppress OIR progression and tumor angiogenesis. Mechanistically, ACE2 Ser-680 phosphorylation and dephosphorylation regulate EC proliferation, sprouting, and migration. The underlying molecular basis is based on phosphorylated ACE2 Ser-680 reducing the activator protein-1 (AP-1)-regulated transcriptomes at the genome-wide scale, thereby inhibiting the transcription of inflammatory genes, which are necessary for angiogenesis.

## Materials and Methods

### Data availability

The data that supports the findings of this study are available from the corresponding author upon request.

Detailed description of the materials and methods is provided in supplementary material. Animal experiments were approved by the Institutional Animal Ethics Committee of Xi’an Jiaotong University (XJTUAE2024-1890).

## Results

### ACE2 Ser-680 Dephosphorylation in Pathological Angiogenesis

Initially, we used the OIR model^18^ to test whether ACE2 Ser-680 phosphorylation status in ECs is involved in the pathological angiogenesis. Levels of ACE2 Ser-680 phosphorylation and total ACE2 were decreased in retinal tissue in the OIR group as compared with sham controls (Figure 1A). With decreased ACE2 Ser-680 phosphorylation in the OIR model, we further investigated the role of Ser-680 phosphorylation/dephosphorylation in angiogenesis *in vivo* by using ACE2D (Arg replacement of Ser-680, phospho-mimetic) and ACE2L (Leu replacement of Ser-680, dephospho-mimetic) mice (Figure 1B). At P17, retinas from ACE2L mice showed a significant increase in neovascular tuft area [assessed by EC marker IsoB4 (in red)]) and decrease in avascular area as compared with ACE2D mice (Figure 1C).

**Figure 1.**
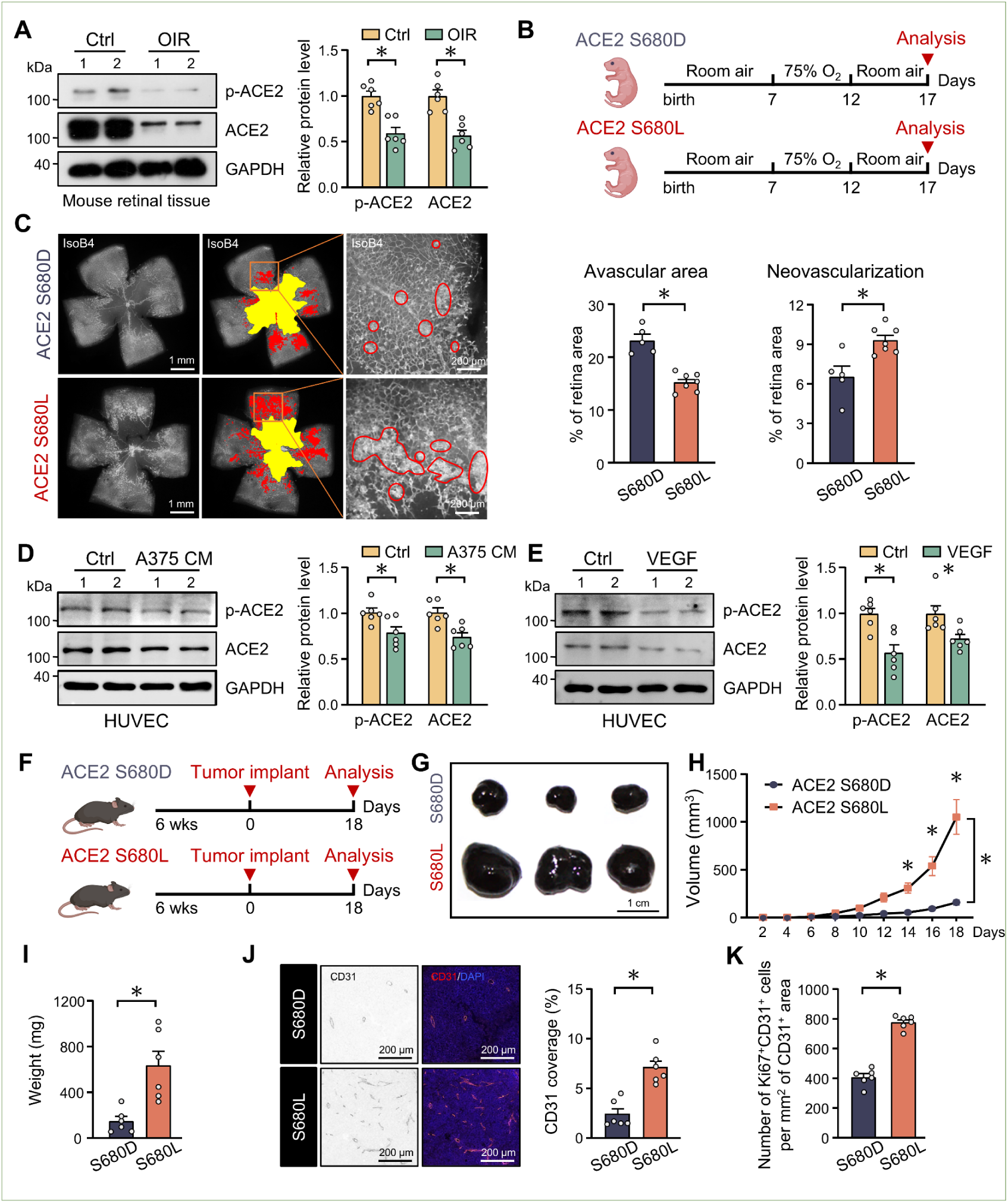
ACE2 Ser-680 Dephosphorylation Is Associated with Pathological Angiogenesis. (**A**) Western blot analysis of levels of p-ACE2 Ser-680 and ACE2 in retina tissues of oxygen-induced retinopathy (OIR) (n=6) and control mice (n=6). (**B**) Schematic diagram illustrating the OIR ACE2 S680D and S680L mouse models. (**C**) Representative images of the isolectin B4-stained retinal vasculature of ACE2 S680D and ACE2 S680L mice under OIR. Areas in yellow indicate the avascular area and those in red neovascular tufts. Scale bar: left, 1 mm; right, 200 µm. Shown on the right are quantifications of avascular area and neovascularization in retina of ACE2 S680D (n=5) and ACE2 S680L (n=7) mice. (**D, E**) Western blot analysis of levels of p-ACE2 Ser-680 and ACE2 in human umbilical vein endothelial cells (HUVECs) cultured with conditioned media from A375 human melanoma cell line (D), vascular endothelial growth factor (VEGF; 50 ng/mL) or PBS (E) for 24 h (n=6, per group). (**F**) Schematic diagram of subcutaneous injection of B16-F10 cells (5×10^5^ cells/mouse) in ACE2 S680D and ACE2 S680L mice (6-8 weeks) to induce melanoma. (**G**) Representative images of explanted B16-F10 tumors from ACE2 S680D and ACE2 S680L mice. Scale bar=1 cm. (**H**) Growth curves for subcutaneous B16-F10 tumors in ACE2 S680D and ACE2 S680L mice. (**I**) End-stage tumor weight in ACE2 S680D and ACE2 S680L mice. (**J**) Representative images of CD31-positive vessels (red) and DAPI nuclear staining (blue) in tumor tissues from ACE2 S680D or ACE2 S680L mice. Scale bar=200 µm. The right panel is quantification of blood vessel density in tumors isolated from ACE2 S680D and ACE2 S680L mice. (**K**) Quantification of endothelial cell (EC) proliferation (Ki67 staining) of tumors from ACE2 S680D and ACE2 S680L mice. For panel G-K, n=6 each group. Data are mean±SEM. \**P* < 0.05. Normally distributed data were analyzed by 2-tailed Student *t* test in (A), (C), (D), (E, pACE2 level) and (K) or Welch *t* test in (I). Non-normally distributed data in (E, ACE2 level) (J) were analyzed by Mann-Whitney *U* test. Two-way ANOVA multiple comparisons with Holm-Šídák post-hoc test among multiple groups in (H).

To further interrogate the role of ACE2 phosphorylation in tumor angiogenesis, we incubated ECs with conditioned media collected from A375 human melanoma cells. ACE2 Ser-680 phosphorylation and total ACE2 level were decreased in ECs (Figure 1D). In addition, treatment with VEGF, a potent angiogenic factor, decreased ACE2 Ser-680 phosphorylation and ACE2 level in ECs (Figure 1E). Given that tumor growth is highly related to angiogenesis,^5^ we next compared tumorigenesis in ACE2D and ACE2L mice xenografted with B16-F10 murine melanoma cells (Figure 1F). As demonstrated by increased volume and weight, tumor growth was more pronounced in male and female ACE2L than ACE2D mice (Figure 1G-I, Supplemental Figure 1A-E). In line with the increased tumorigenesis, CD31 staining and EC proliferation were more profound in xenografted tumors from ACE2L than ACE2D mice (Figure 1J-K). The pro-angiogenic nature in ACE2L mice was further validated by using LLC cells for xenografting (Supplemental Figure 1F-J). Collectively, the results in Figure 1 suggest that ACE2 Ser-680 dephosphorylation in ECs is indispensable in pathological angiogenesis.

### ACE2 Ser-680 Dephosphorylation Is Pro-angiogenic *In Vitro*

ECs involved in pathological angiogenesis show increased glycolysis, proliferation, and migration.^19^ Having previously shown that ACE2 Ser-680 dephosphorylation increased glycolysis in ECs,^20^ we analyzed whether EC proliferation and migration are increased by ACE2 Ser-680 dephosphorylation. As compared with adenovirus-mediated overexpression of ACE2 S680D in ECs, that of ACE2 S680L increased the EC population in the S phase (Figure 2A). The increased proliferation by ACE2 S680L was verified by EdU incorporation assay (Figure 2B). In line with these results, ACE2 S680L overexpression increased spheroid sprouting (Figure 2C) and tube formation (Figure 2D). The effect of ACE2 Ser-680 dephosphorylation on microvascular sprouting was examined by aortic ring angiogenesis assay *ex vivo*. The formation of microvascular networks surrounding the aortic ring was more prominent in ACE2L than ACE2D mice (Figure 2E). During angiogenesis, tip ECs have long and dynamic filopodia.^21^ ECs overexpressing ACE2 S680L or stimulated with VEGF exhibited phalloidin-positive filopodia, which were fewer in ECs overexpressing ACE2 S680D (Figure 2F). Thus, ECs with dephosphorylated ACE2 Ser-680 tended to be pro-angiogenic *in vitro*.

**Figure 2.**
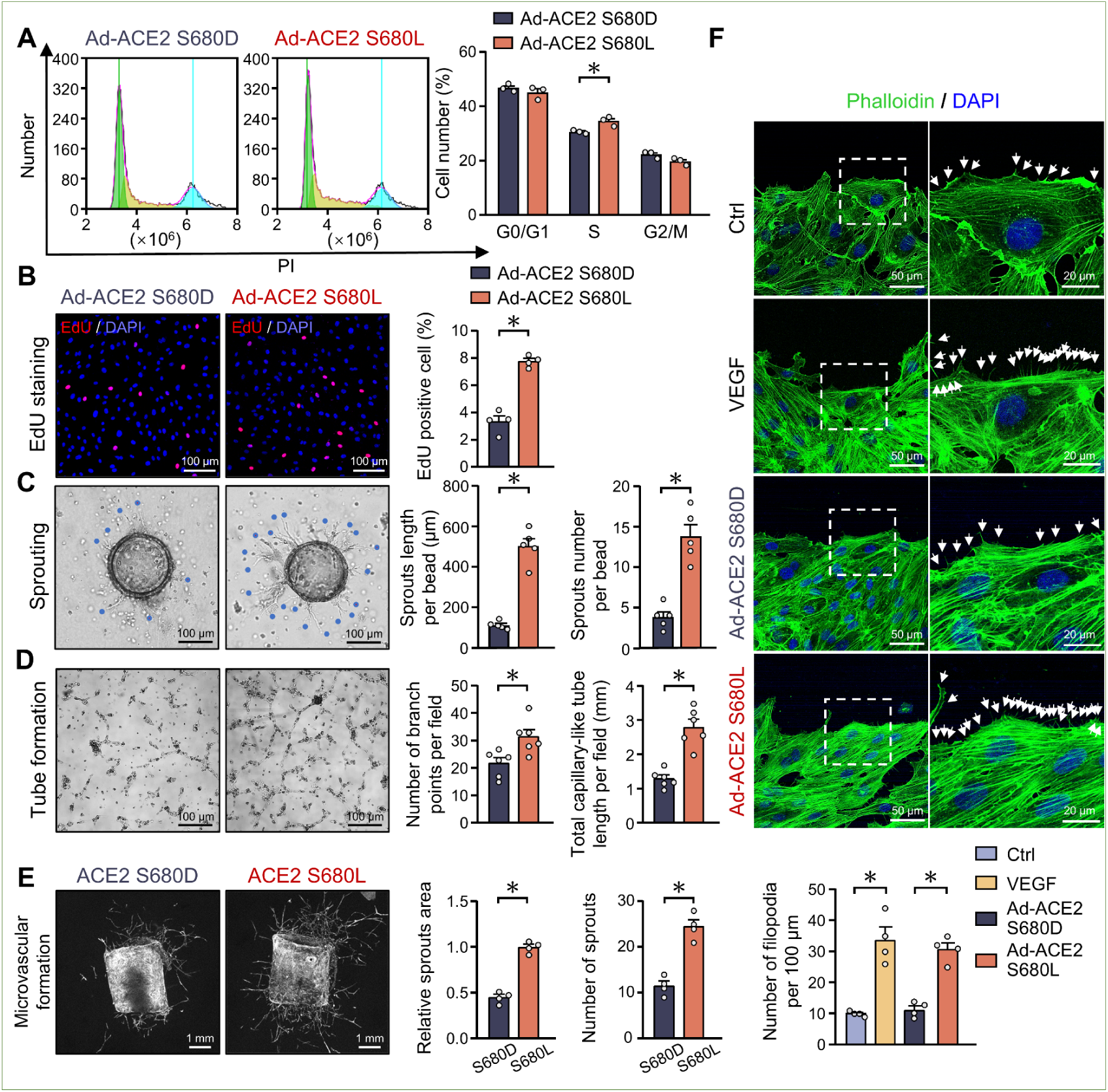
ACE2 Ser-680 Dephosphorylation Promotes Angiogenesis *In Vitro*. (**A-D**) HUVECs were infected with Ad-ACE2 S680D or Ad-ACE2 S680L (10 multiplicity of infection [MOI]) for 48 h. (**A**) EC cell cycle progression was analyzed by flow cytometry. (**B**) EdU immunostaining (red) for EC proliferation assay, with nuclei counterstained by DAPI (blue). Scale bar=100 µm. (**C**) Representative images and quantification of endothelial sprouting in fibrin gel. Scale bar=100 µm. (**D**) Representative image of tube formation. The visible tubes in randomly selected fields were determined by number of branch points and total tube length. Scale bar=100 µm. (**E**) EC sprouts protruding from aortic rings isolated from ACE2 S680D and ACE2 S680L mice (6-8 weeks, n=4 each group). Representative images were taken at day 5 of cultivation. Scale bar=1 mm. (**F**) Filopodia formation assays of HUVECs overexpressing ACE2 S680D or ACE2 S680L or stimulated with VEGF (50 ng/mL) or PBS. Filopodia, revealed by phalloidin staining (white arrows), at the leading edge were imaged 6 h after scratching. Scale bars=50 µm. Insets are 3X magnified. Scale bars=20 µm. Data are mean±SEM. \**P* < 0.05. Normally distributed data were analyzed by 2-tailed Student *t* test in (A-E). Two-way ANOVA test with Dunnett’s multiple comparisons test among multiple groups in (F).

### ACE2 Ser-680 Dephosphorylation Activates Angiogenic Pathways

Given the EC angiogenic phenotype driven by ACE2 Ser-680 dephosphorylation, we then examined the underlying transcriptomic changes in ECs. Differentially expressed gene (DEG) analysis (log2FC>0.5 and FDR<0.05) revealed that a total of 1998 genes were upregulated by ACE2 S680L versus ACE2 S680D (Figure 3A), and a significant number of these were involved in angiogenesis-related pathways, including cell proliferation, migration, cell cycle, and the VEGF signaling pathway, as demonstrated by KEGG plots in Figure 3B. Heatmaps in Figure 3C show that these DEGs were related to proliferation and migration, proinflammatory genes, and pro-angiogenic genes. Furthermore, gene set enrichment analysis demonstrated enrichment of the VEGF signaling pathway by ACE2 S680L (Figure 3D). RT-qPCR validated that ACE2 S680L upregulated transcripts encoding *CXCL8*, *CCL2*, *IL6* and *NFKB1* (pro-inflammation); *VEGFA*, *ANGPT2* and *ROCK1* (pro-angiogenesis); and *MAPK1*, *S1PR1* and *MAP2K1* (pro-migration) (Figure 3E). As controls, *KLF4* and *NOS3* were downregulated (Figure 3E). We further studied upregulation of the angiogenic genes at the translational level by using a Proteome Profiler mouse angiogenesis array. The levels of 33 of 53 angiogenesis-related proteins were higher in OIR retina tissues from ACE2L than ACE2D mice. Especially Cyr61, DLL4, DPP4, Endostatin, FGF-1, IGFBP-1, CXCL10, MCP-1, PDGF-AA, PAI-1, VEGF-A and VEGF-B were highlighted (Figure 3F).

**Figure 3.**
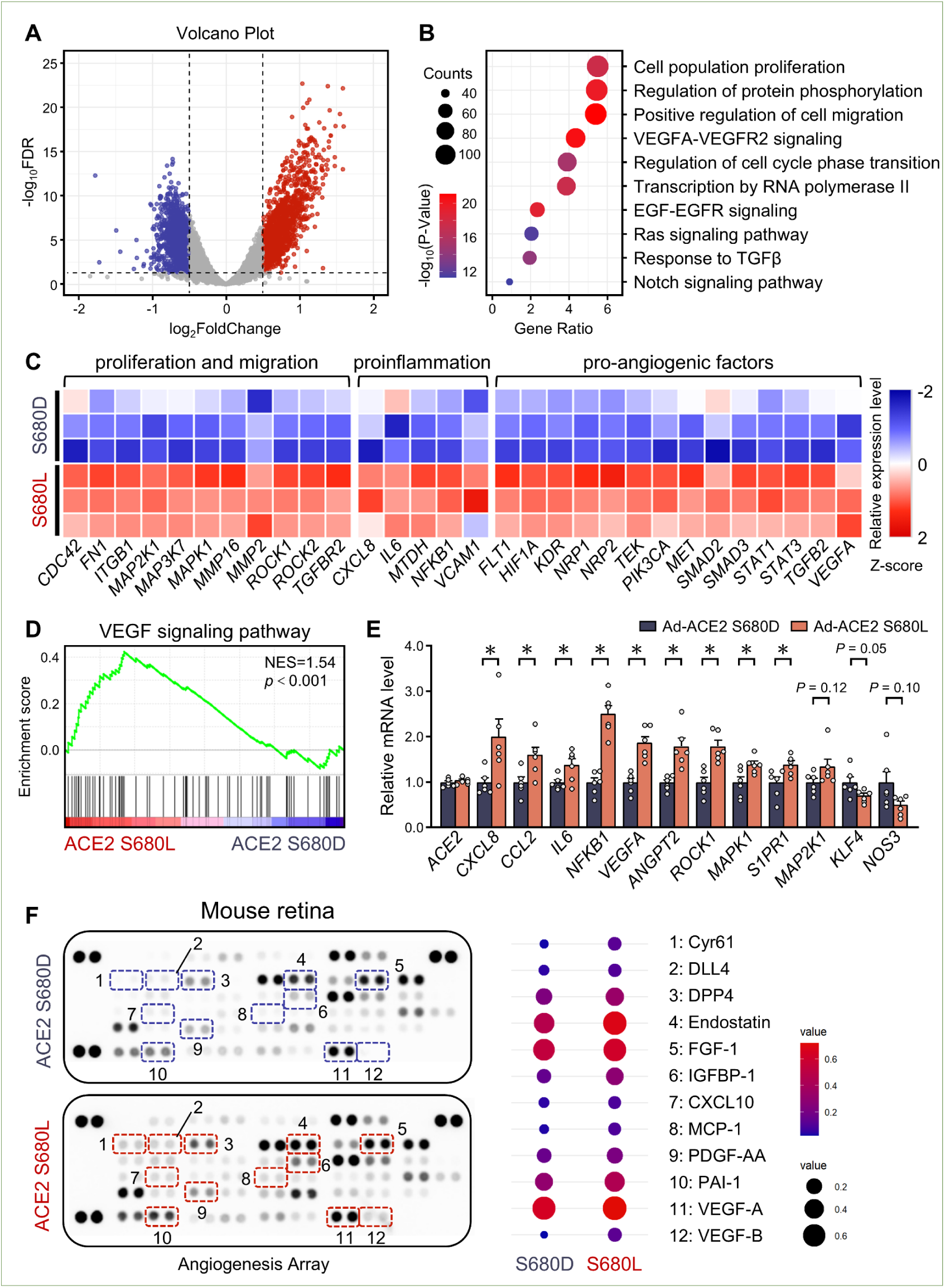
ACE2 Ser-680 Dephosphorylation Activates Pro-angiogenic Genes. (**A-D**) HUVECs were infected with Ad-ACE2 S680D or Ad-ACE2 S680L (10 MOI) for 48 h. Total RNA was isolated for bulk RNA-seq analysis. (**A**) Volcano plot showing transcriptomic changes between ACE2 S680D and ACE2 S680L overexpression (|log2FC|>0.5 and false discovery rate [FDR]<0.05). (**B**) Functional enrichment analysis with DAVID (https://davidbioinformatics.nih.gov) for 1,998 DEGs upregulated by ACE2 S680L, visualized as -log(P value). (**C**) Heatmap of expression of angiogenesis-related DEGs between ACE2 S680D and ACE2 S680L. The relative expression is presented as Z-scores. (**D**) Gene set enrichment analysis (GSEA) of ACE2 S680D versus ACE2 S680L for bulk RNA-seq with VEGF signaling pathway. NES: normalized enrichment score. (**E**) qPCR analysis of mRNA levels of angiogenic genes (n=6). (**F**) Angiogenesis array assay of OIR retina from ACE2 S680D and ACE2 S680L mice. Fold change analysis of differentially expressed angiogenic factors (right panel). Data are mean±SEM. \**P* < 0.05. Normally distributed data were analyzed by 2-tailed Student *t* test in (E, *CCL2*, *IL6*, *NFKB1*, *VEGFA*, *ROCK1*, *MAPK1*, *S1PR1*, *KLF4*) or Welch *t* test in (E, *CXCL8*, *ANGPT2*, *NOS3*). Non-normally distributed data in (E, *MAP2K1*) were analyzed by Mann-Whitney *U* test.

### ACE2 Ser-680 Dephosphorylation Induces Angiogenesis via AP-1

Because ACE2 S680D and S680L differentially regulated gene expression at the global scale, we theorized that ACE2 Ser-680 phosphorylation/dephosphorylation might be involved in the epitranscriptomic and/or transcriptional regulatory mechanisms. This thesis was supported in part by the upregulation of MED1, HDAC9, BTAF1, and BACH1 by ACE2 S680L (Supplemental Figure 2A, 2B). To further verify this, we conducted ATAC-seq analysis of ECs overexpressing ACE2 S680L or ACE2 S680D. Although the chromatin distributions (i.e., promoter, 5’UTR, 3’UTR, 1^st^ exon, etc.) of ATAC signals were approximately similar with the treatments (Figure 4Ai), ACE2 S680L caused an ATAC enrichment at the transcription start site±3 kb region by 4.8% (Figure 4Aii), so ACE2 S680L decondensed the chromatin more than did ACE2 S680D. Leveraging the ATAC-seq data, we analyzed the chromatin regions and the cognate transcription factors (TFs) affected by ACE2 S680L and ACE2 S680D, respectively (Figure 4B). Unsupervised motif-enrichment analysis showed that binding sites corresponding to the AP-1 family of TFs (e.g., JUNB, JUND, JUN, FOSL2 and FOS) were the most enriched in ECs overexpressing ACE2 S680L. However, binding sites for THAP1, NRF1, MYB, CTCF, and CTCFL were preferentially enriched by ACE2 S680D versus ACE2 S680L (Figure 4C). The exemplified ACE2 S680L-enhanced JunB and ACE2 S680D-enhanced CTCF motifs are depicted in Figure 4D. Function-wise, KEGG pathway enrichment analysis of the ATAC-seq data showed that genes with enriched JunB motif were largely associated with pathways in cancer, MAPK, PI3K-Akt, HIF-1, Hippo signaling pathways which were highly associated with angiogenesis (Supplemental Figure 2C). Henceforth, we focused on JUNB as a representative member of the AP-1 family of TFs to substantiate how ACE2 Ser-680 dephosphorylation induces angiogenic genes via AP-1 induction. In the following validation experiments, ACE2 S680L overexpression increased the transcriptional upregulation of *JUNB*, *JUND*, *JUN*, *FOS* and *FOSL2* (qPCR in Figure 4E), AP-1 binding to the TGACTCA cognate motif (EMSA assay in Figure 4F), and JUNB nuclear sequestration (immunostaining in Figure 4G). Correspondingly, a luciferase reporter driven by four copies of AP-1 element (4×TRE-Luc)^22^ was significantly induced by ACE2 S680L versus ACE2 S680D overexpression (Figure 4H). As a control, oxPAPC, an angiogenic and inflammatory stimulator^23^ induced 4×TRE-Luc. Collectively, results in Figure 4 suggested that angiogenic cues (e.g., cancer microenvironment, inflammation, and hypoxia) might induce angiogenic genes via ACE2 Ser-680 dephosphorylation, which promoted the AP-1 (e.g., JUNB)-mediated transactivation.

**Figure 4.**
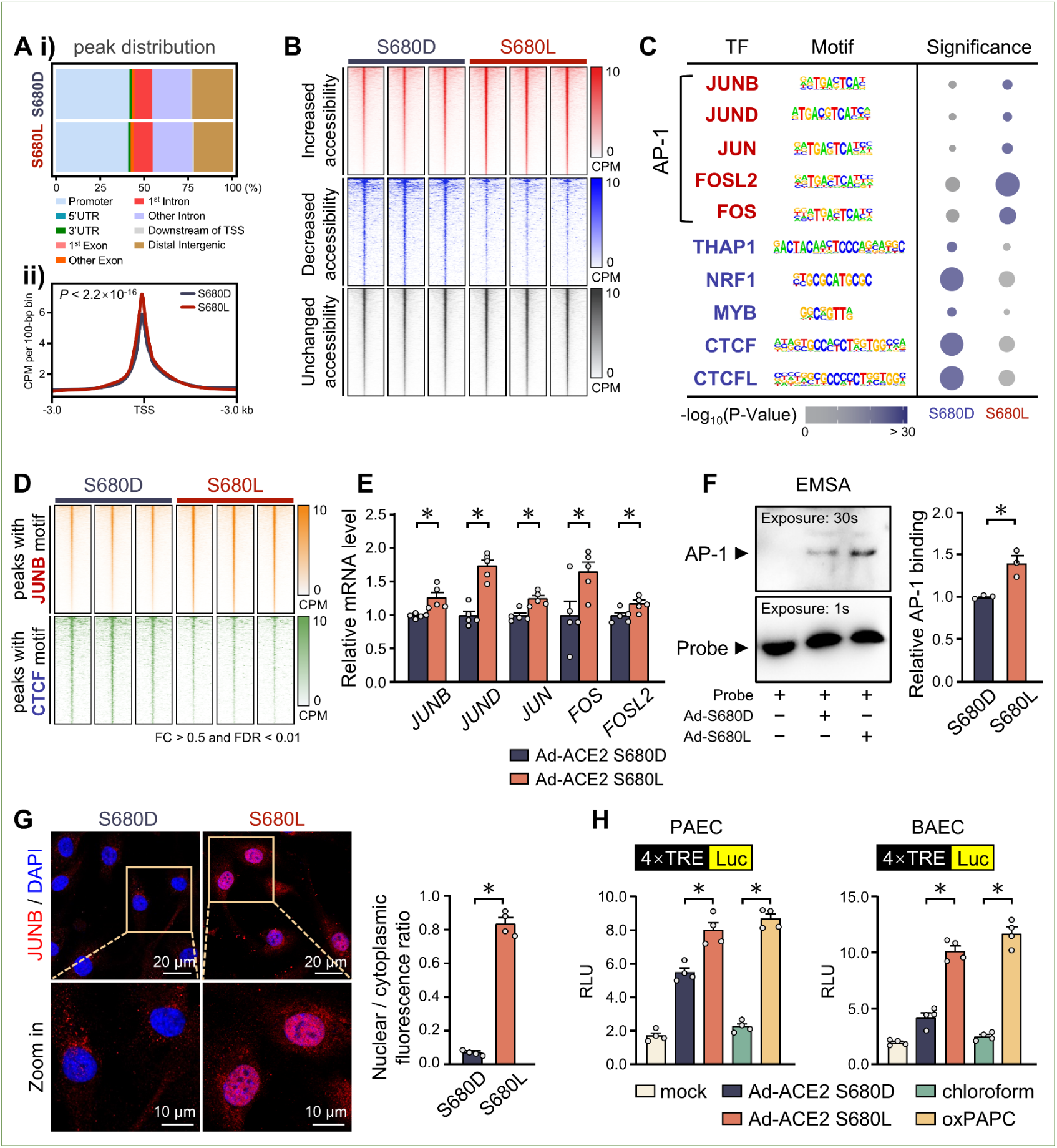
ACE2 Dephosphorylation Increases Chromatin Accessibility of AP-1. HUVECs overexpressing ACE2 S680D or ACE2 S680L underwent ATAC-seq analysis. (**Ai**) Annotation of genome-wide chromatin accessible regions identified by ATAC-seq. UTR, untranslated region; TSS, transcription start site. (**Aii**) Aggregated ATAC-seq signals across the TSS±3 kb region showing reduced chromatin accessibility in ECs overexpressing ACE2 S680D versus ACE2 S680L. CPM, counts per million. (**B**) ATAC-seq footprinting heatmap showing differential accessibility between ACE2 S680D and ACE2 S680L (FDR < 0.1). (**C**) Motif-enrichment analysis indicating AP-1 family transcription factor upregulation by ACE2 S680L and THAP1, NRF1, MYB, CTCF, and CTCFL upregulation by ACE2 S680D. (**D**) ATAC-seq footprinting heatmap showing differential accessibility by ACE2 S680D versus ACE2 S680L at *loci* of JUNB- or CTCF-corresponding motifs (FC > 0.5 and FDR < 0.01). (**E**) qPCR analysis of mRNA levels of AP-1 family genes in ECs overexpressing ACE2 S680D or ACE2 S680L. (**F**) EMSA revealing increased binding of AP-1 to the TRE (i.e., TGACTCA) by ACE2 S680L overexpression. (**G**) Representative immunostaining of JUNB nuclear sequestration in ECs overexpressing ACE2 S680L. Top: Scale bar=20 µm, Bottom: Scale bar=10 µm. (**H**) Pulmonary artery ECs and bovine artery ECs were transfected with a luciferase reporter fused with 4×TRE, then treated with Ad-ACE2 S680D or Ad-ACE2 S680L for 48 h and chloroform or ox-PAPC (30 μg/mL) for 4 h. Luciferase activity was measured accordingly. Data are mean ±SEM. \**P* < 0.05. Normally distributed data were analyzed by 2-tailed Student *t* test in (E, *JUND*, *JUN*, *FOS*, *FOSL2*) or Welch *t* test in (E, *JUNB*), (F), (G). Two-way ANOVA test with Dunnett’s multiple comparisons test among multiple groups in (H).

Mining data from RNA-seq (Figure 3), ATAC-seq (Figure 4), and JUNB ChIP-seq (GSE89970), we found that the JUNB ChIP-seq signal was commonly enhanced in the promoter region of 502 genes upregulated by ACE2 S680L (Figure 5A). These genes are functionally related to positive regulation of angiogenesis (GO:0045766), VEGF signaling pathway (GO:0038084), cell migration (GO:0030334), EC proliferation (GO:0001935), and cellular response to angiotensin (GO:1904385) (Figure 5B). In confirmation, ACE2 S680L caused a less condensed promoter region of the angiogenesis-related genes (e.g., *KDR*, *FLT1*, *MET* and *TEK*), as revealed by the enhanced ATAC signal (Figure 5Ci-iv). Next, ChIP-qPCR was used to validate that ACE2 S680L overexpression indeed enhanced the binding of JUNB to the promoters of *KDR*, *FLT1*, *MET* and *TEK* (Figure 5D). SiRNA-mediated JUNB knockdown reduced the ACE2 S680L-induced *KDR*, *FLT1*, *MET* and *TEK* transcriptional and translational levels (Figure 5E, Supplemental Figure 3A). Phenotypically, JUNB knockdown in ECs alleviated ACE2 S680L-promoted proliferation, sprouting, migration, and tube formation (Figure 5F). Together, results in Figure 5 suggest that ACE2 Ser-680 dephosphorylation induced an angiogenic phenotype of ECs via JUNB/AP-1–mediated transactivation of angiogenic genes.

**Figure 5.**
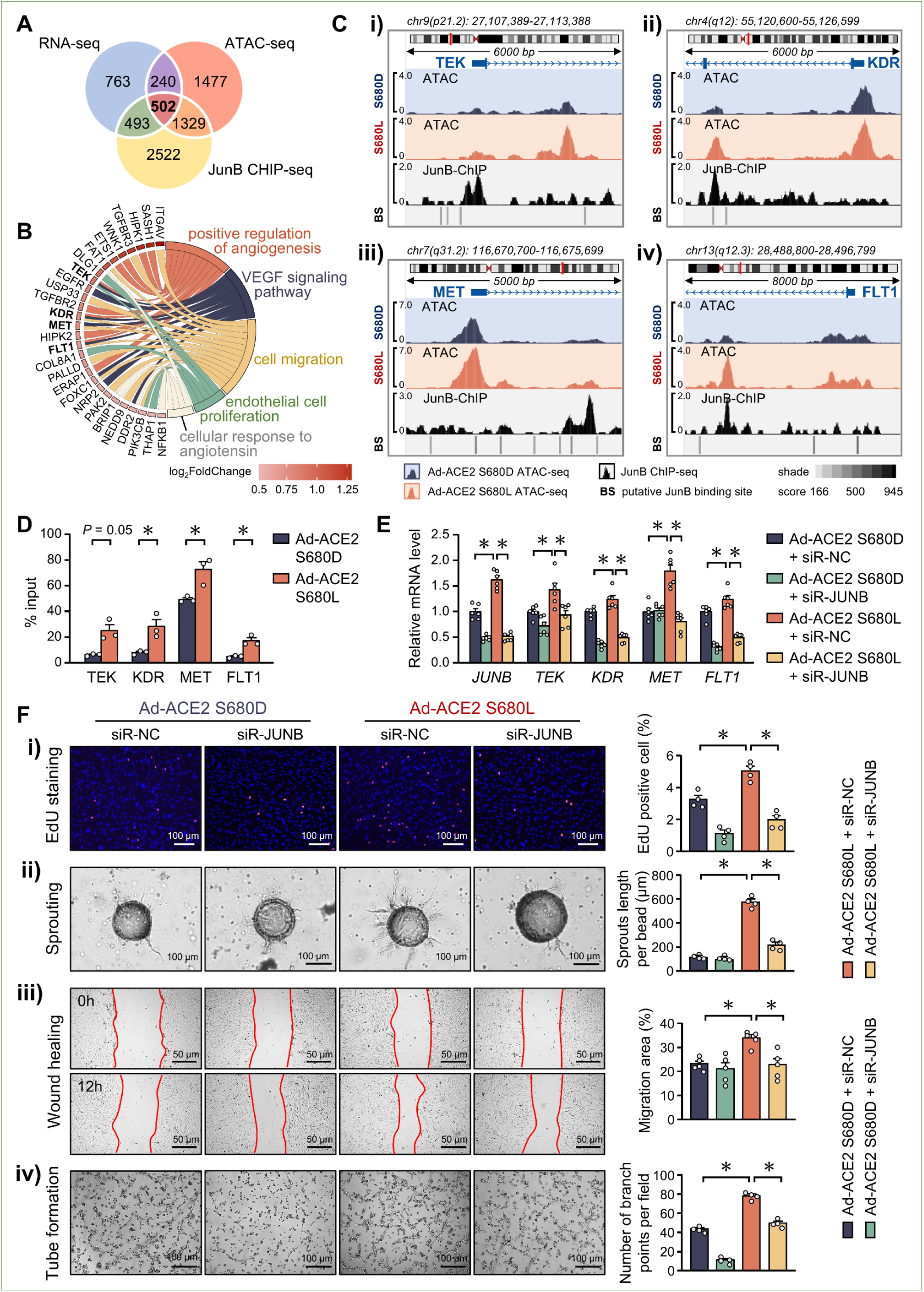
ACE2 Dephosphorylation Induces Pro-angiogenic Genes via AP-1 Transactivation. (**A**) Venn diagram showing the overlap among ACE2L-upregulated genes (n = 1998), ATAC-seq peaks enhanced by ACE2L (n = 3548), and genes with JUNB binding sites in their promoters (n = 4846). (**B**) Circle plots showing angiogenesis-related pathways enriched among the overlapped genes in (A). (**C**) The ATAC-seq and JUNB ChIP-seq tracks at the promoters of *TEK* (Ci), *KDR* (Cii), *MET* (Ciii) and *FLT1* (Civ) in ECs overexpressing ACE2 S680D and S680L. (**D**) JunB ChIP-qPCR revealing enhanced JUNB binding to the *TEK*, *KDR*, *MET,* and *FLT1* promoters in ECs overexpressing ACE2 S680L. (**E**) HUVECs were transfected with JUNB siRNA or control siRNA, then underwent Ad-ACE2 S680D and Ad-ACE2 S680L infection. The mRNA levels of *JUNB*, *TEK*, *KDR*, *MET* and *FLT1* were measured by qPCR (n=6) after 48 h. (**F**) HUVECs overexpressing ACE2 S680D or S680L with or without JUNB knockdown were used for EdU proliferation, Scale bar=100 µm (Fi), spheroid sprouting, Scale bar=100 µm (Fii), wound healing, Scale bar=50 µm (Fiii), and tube formation, Scale bar=100 µm (Fiv) assays. Data are mean±SEM. \**P* < 0.05. Normally distributed data were analyzed by 2-tailed Student *t* test in (D, *KDR*, *FLT1*, *MET*) or Welch *t* test in (D, *TEK*). Two-way ANOVA test with Dunnett’s multiple comparisons test among multiple groups in (E) and (F).

### Aggravated Angiogenesis in ACE2L Mice

To ensure that the ACE2 Ser-680 dephosphorylation–enhanced angiogenesis is mediated by JUNB/AP-1 *in vivo*, ACE2L and ACE2D mice underwent OIR and were administered T-5224, a selective AP-1 inhibitor, or not (Figure 6A).^24^ T-5224 inhibition of JUNB/AP-1 in the retina of ACE2L mice was confirmed by decreased JUNB phosphorylation (Supplemental Figure 3B), and *Kdr*, *Flt1*, *Met*, and *Tek* levels were reduced at the transcriptional level (Supplemental Figure 3C). Consequent to this AP-1 inhibition, angiogenesis was mitigated, as indicated by the reduced neovascularization area in ACE2L mice (Figure 6B-C).

**Figure 6.**
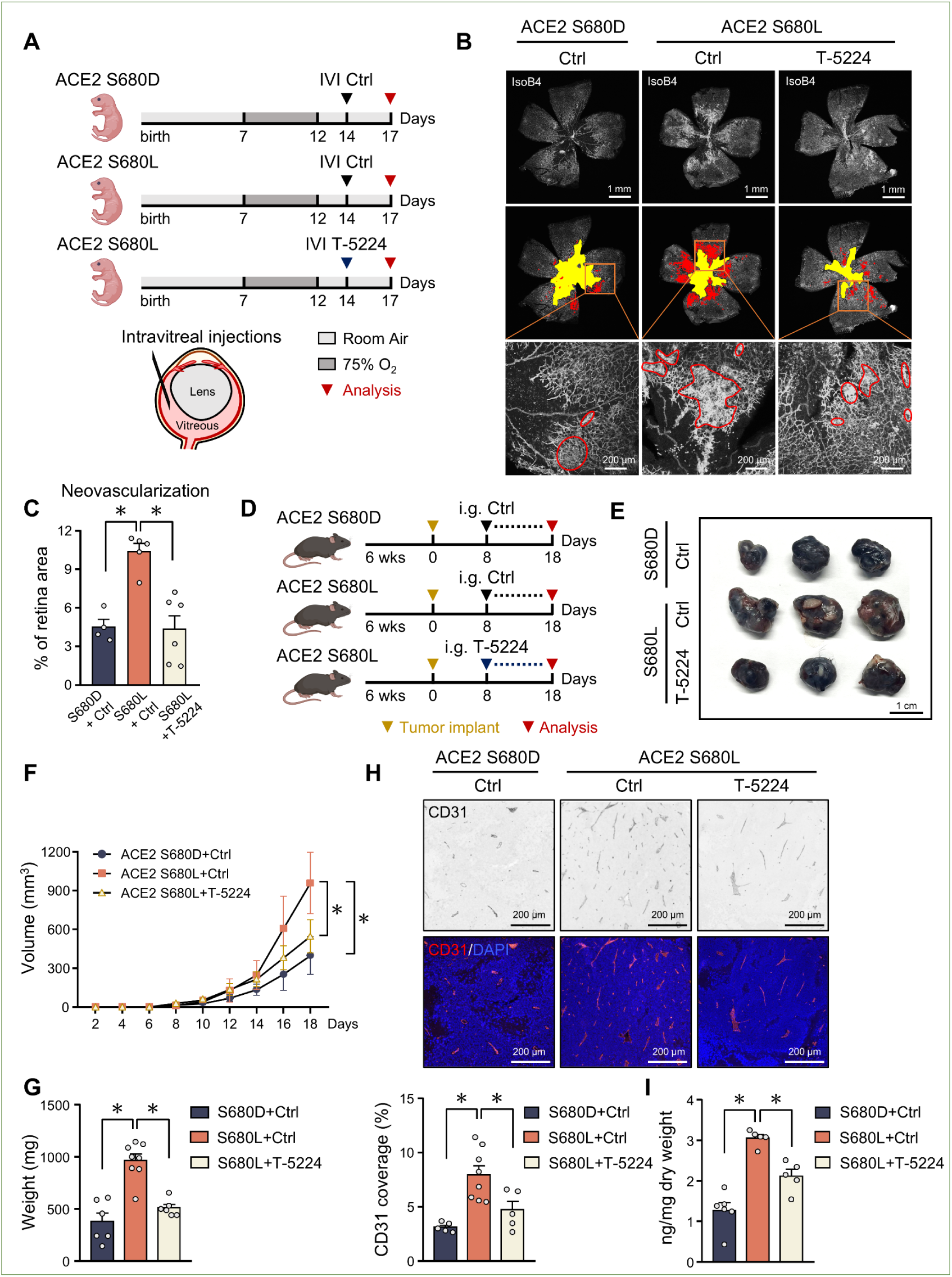
ACE2 S680L-AP-1 Axis Aggravates Angiogenesis *In Vivo*. (**A**) Schematic diagram illustrating the OIR mouse models involving AP-1 inhibition. (**B**) Representative images of the isolectin B4-stained retinal vasculature of ACE2 S680D and ACE2 S680L mice with or without administration of AP-1 inhibitor T-5224. Areas in yellow indicate the avascular area and those in red neovascular tufts. Top: Scale bar=1 mm, Bottom: Scale bar=200 µm. (**C**) Quantification of neovascularization area in retina of various mouse groups. (**D**) Schematic diagram illustrating the B16-F10 tumor mouse model with or without T-5224 inhibition of AP-1. (**E**) Representative images of explanted B16-F10 tumors in the 3 groups of mice; ACE2 S680D mice and ACE2 S680L mice with or without T-5224 treatment. Scale bar=1 cm. (**F**) Growth curves for subcutaneous B16-F10 tumors, (**G**) End-stage tumor weight, (**H**) Representative images of blood vessels in tumors stained with CD31 (red) and DAPI (blue), Scale bar=200 µm. (**I**) Vascular permeability revealed by Evans blue extravasation in the 3 groups of mice in (D). Data are mean±SEM from 4-8 mice per group. *P <0.05. Two-way ANOVA test with Dunnett’s multiple comparisons test among multiple groups in (C), (G), (H) and (I). Two-way ANOVA multiple comparisons with Holm-Šídák post-hoc test among multiple groups in (F).

ACE2L mice administered T-5224 were also used for the melanoma angiogenesis model (Figure 6D). Tumor volume and weight were reduced in ACE2L mice administered T-5224 as compared with mice receiving vehicles (Figure 6E-G). CD31 immunostaining and Evans blue staining recapitulated that T-5224 inhibition of the JUNB/AP-1–reduced vascular density (Figure 6H) and improved vascular integrity (Figure 6I). These experiments involving ACE2L mice and T-5224 validated the causality among ACE2 S680 dephosphorylation, JUNB/AP-1 activation, and pathogenic angiogenesis *in vivo*.

### ACE2 Lys-788 Deubiquitination Is Anti-angiogenic

ACE2 S680 dephosphorylation promotes the K48-linked ACE2 ubiquitination at Lys-788 and thereafter ACE2 degradation.^17^ With the increased ACE2 ubiquitination in the retinal tissues of wild-type (WT) mice undergoing OIR (Supplemental Figure 4A), we tested whether deubiquitination of ACE2 at K788 produces an anti-angiogenic phenotype. Overexpression of ACE2 K788R (deubiquitination-mimetic mutant) in cultured ECs attenuated VEGF-induced filopodia protrusion at the leading edge and JUNB nuclear sequestration (Figure 7A), in line with AP-1 attenuation. Furthermore, ACE2 K788R or S680D overexpression reduced the VEGF-induced ACE2 ubiquitination (Supplemental Figure 4B). To this end, we created an ACE2 K788R knock-in mouse line. Analysis of the retinal vasculature at P17 showed that pathological neovascularization was strongly reduced in ACE2 K788R mice compared with their WT littermates (Figure 7B). These mice also showed decreased numbers of filopodia and tip cells in P6, which phenocopied that in ACE2D mice (Supplemental Figure 5). As well, ACE2 K788R mice showed reduced tumor size and reduced angiogenesis in the melanoma xenograft model as compared with their WT littermates (Figure 7C-F). These results suggest that ACE2 Ser-680 phosphorylation/dephosphorylation, rendering Lys-788 deubiquitination/ubiquitination, determined the angiogenic phenotype.

**Figure 7.**
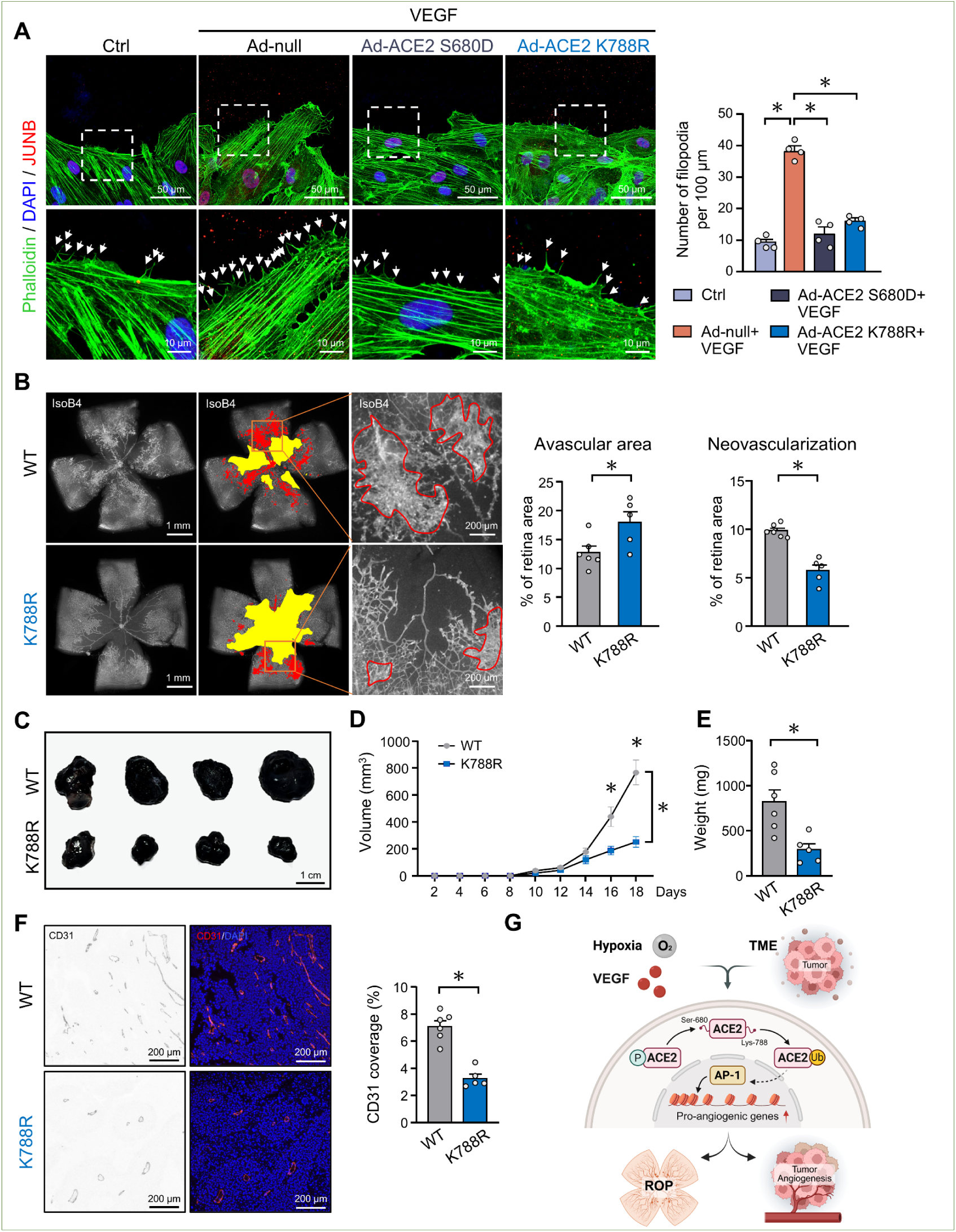
ACE2 Lys-788 Deubiquitination Phenocopies the Anti-angiogenic ACE2 Ser-680 Phosphorylation. (**A**) Filopodia assessment of HUVECs infected with Ad-null, Ad-ACE2 S680D, or Ad-ACE2 K788R and stimulated with VEGF (50 ng/mL). Filopodia at the leading edge, revealed by phalloidin staining (white arrows), were imaged 6 h after scratching. Scale bars=50 µm. Insets are 3X magnified. Scale bars=10 µm. (**B**) Representative images of the isolectin B4-stained retinal vasculature of ACE2 K788R and WT mice under OIR. Areas in yellow indicate the avascular area and those in red neovascular tufts. Scale bar: left, 1 mm; right, 200 µm. Shown on the right are quantifications of avascular area and neovascularization in retina of ACE2 K788R (n=5) and WT (n=6) mice. (**C**) Representative images of explanted B16-F10 tumors in ACE2 K788R and WT littermate mice. Scale bar=1 cm. (**D**) Growth curves of subcutaneous B16-F10 tumors in ACE2 K788R and WT mice. (**E**) End-stage tumor weight in ACE2 K788R and WT mice. (**F**) Representative images of blood vessels in ACE2 K788R and WT tumors stained with CD31 (red) and DAPI (blue). Scale bar=200 µm. On the right is the quantification of blood vessel density. In panel C-F, n=5-6 mice in each group. Data are mean±SEM. \**P* < 0.05. Normally distributed data were analyzed by 2-tailed Student *t* test in (E), (F). Two-way ANOVA test with Dunnett’s multiple comparisons test among multiple groups in (A), (B). Two-way ANOVA multiple comparisons with Holm-Šídák post-hoc test among multiple groups in (D). (**G**) The proposed working model illustrating ACE2 Ser-680 dephosphorylation regulation of angiogenesis via AP-1. Hypoxia, tumor microenvironment, or VEGF stimulation leads to dephosphorylated ACE2 Ser-680, which promotes Lys-788 ubiquitination. This mode of ACE2 modification in ECs leads to AP-1–regulated expression of the pro-angiogenic genes.

## Discussion

In this study, we demonstrated that ACE2 Ser-680 phosphorylation/dephosphorylation in ECs is indispensable in angiogenesis. The following observations support this thesis: (1) ACE2 Ser-680 phosphorylation was reduced in OIR tissues from mice with retinal angiogenesis and tumor ECs; (2) ACE2 S680L mice exhibited exacerbated pathological angiogenesis, including both OIR and tumor angiogenesis; (3) ACE2 S680L ECs showed increased proliferation, migration and sprouting; and (4) ACE2 K788R phenocopied the anti-angiogenic phenotype of ACE2 S680D. Mechanistically, as summarized in Figure 7G, ACE2 Ser-680 dephosphorylation/Lys-788 ubiquitination enhanced chromatin accessibility for the AP-1 family of transcription factors, thereby increasing the expression of angiogenic genes.

Regulating blood pressure, fluid balance, and electrolyte homeostasis, RAS is balanced between the detrimental ACE1-Ang II-AT1R axis and the protective ACE2-Ang-(1–7)-Mas axis. Ang II acting on AT1R induces AP-1– and NFκB-dependent gene expression. An imbalanced RAS in the vascular tissue predisposes to endothelial dysfunction, which contributes substantially to various cardiovascular diseases.^25^ Dysregulated RAS is involved in the pathogenesis of ROP, because elevated levels of RAS component proteins and peptides are present in the vitreous of patients with ROP^10^ and gain-of-function variants of ACE1 are associated with ROP progression and treatment outcomes.^26^ Also, pharmacologic interventions of RAS could inhibit ROP progression and preserve the retina.^10^ In tumor angiogenesis, expression of AT1R is increased in many types of carcinomas. RAS inhibitors (ACEI or ARB) reduce the risk of cancers in the digestive, respiratory, urologic, gynecologic, and dermatologic systems,^27–31^ which may be due in part to the rebalance of RAS.

Here, we used ROP and skin cutaneous melanoma as two disease models to elucidate the causality between ACE2 PTMs and angiogenesis. In line with decreased ACE2 Ser-680 phosphorylation found in both models (Figure 1), phosphorylated Ser-680 (mimicked by S680D) and deubiquitinated Lys-788 (mimicked by K788R) mitigated angiogenesis in ROP and skin cutaneous melanoma (Figures 1, 7). Recently, ACE2 deficiency was found to promote hepatocellular carcinoma (HCC) development and anti-PD-L1 resistance, but ACE2 overexpression inhibited HCC progression in immune-competent mice.^32^ Additionally, ACE2 may also facilitate tumor antigen presentation and tumor killing.^33^ Thus, ACE2 Ser-680 dephosphorylation may also participate in these tumorous activities.

AMPK plays a context-dependent role in angiogenesis.^34^ In the energy-demanding state resulting from hypoxia, ischemia, and exercise, AMPK is activated and has protective effects, including promoting angiogenesis.^35–37^ AMPK is inhibited under pathologic conditions, such as retinopathy and cancer. Thus, sustained AMPK activation often attenuates pathologic angiogenesis,^38,39^ and AMPK activators, such as resveratrol, curcumin, and epigallocatechin gallate, exhibit anti-angiogenic properties.^40–42^ AMPK phosphorylates ACE2 Ser-680, which is consistent with the anti-angiogenic effect of ACE2 S680D.^16^ Given that metformin has an inhibitory effect on both OIR and tumor angiogenesis,^43,44^ we theorize that this effect is mediated in part via AMPK phosphorylation of ACE2 Ser-680.

Dysregulated angiogenesis has been documented in multiple organs in people with COVID-19.^45,46^ A hyperinflammatory state and upregulated angiogenic genes were also found in various organs in SARS-CoV-2–infected rhesus macaques.^47^ Intussusceptive angiogenesis seems to be a feature of COVID-19 pathology.^48^ For example, in lungs from deceased COVID-19 patients, the abrupt intussusceptive angiogenesis was associated with upregulation of 59 angiogenesis-related genes.^45^ Intriguingly, most of these proliferation-, migration-, and inflammation-related genes were also upregulated in ECs overexpressing ACE2 S680L versus those expressing S680D in our study (Supplemental Figure 5). We previously demonstrated that the SARS-CoV-2 spike protein dephosphorylates ACE2 Ser-680, which leads to increased glycolysis and decreased oxidative phosphorylation in ECs.^20^ These metabolic switches, like the Warburg effect, facilitate pillar formation and intussusceptive remodeling.^49,50^ Using COVID-19 vasculitis as an example, we propose that PTMs of ACE2 may participate in various forms of pathogenic angiogenesis, whose universality and clinical implications warrant further study.

ACE2 is a membrane-bound metallopeptidase; Ser-680 resides in the a.a 18-740 extracellular domain and Lys-788 in the a.a 762-805 cytoplasmic domain (UniProt Q9BYF1).^51^ The subcellular locations where Ser-680 and Lys-788 are modified and how they coordinately regulate the epigenetics and transcription remain unclear. Recent studies suggest that the C-terminal tail of ACE2 contains a highly conserved nuclear localization signal and that the nuclear-compartmented ACE2 colocalizes with RNA polymerase II.^52,53^ Thus, like several angiogenesis-related receptors, such as VEGF receptor and epidermal growth factor receptor,^54–56^ ACE2 may also have nuclear functions mediated by PTMs. If so, under certain pro-angiogenic conditions, such as hypoxia and VEGF stimulation, dephosphorylated/ubiquitinated ACE2 likely translocates to the nucleus to regulate genes engaged in proliferation, migration, inflammation, and angiogenesis (Figure 3).

The AP-1 family of dimeric transcription complexes plays a pivotal role in a broad spectrum of vascular pathologies such as aortic dissection, atherosclerosis, and vascular aging.^24,57,58^ VEGF induces EC migration and proliferation via AP-1 activation, which represents a key angiogenic mechanism involved in multiple tumor cell lines and ECs.^59^ Here, we showed that dephosphorylated/ubiquitinated ACE2 affected chromatin remodeling and transcriptional activation of AP-1–regulated genes, conducive to pathological angiogenesis. (Figure 4). As the selective AP-1 inhibitor, T-5224 specifically inhibits the DNA binding activity of c-Fos/c-Jun at the AP-1 regulatory element. With its inhibition of AP-1–regulated inflammatory cytokines, T-5224 is used in clinical studies for atopic dermatitis, acute kidney injury, and head and neck squamous cell carcinoma.^60–62^ Our findings that local or systemic delivery of T-5224 inhibited OIR or melanoma angiogenesis are consistent with findings that EC-specific JUNB knockout impairs expression of neurovascular guidance genes and disrupts the formation of the retinal vascular plexus.^63^ Of note, sphingosine 1-phosphate receptor (S1PR)-elicited signaling induces this developmental angiogenesis via JUNB.^63^ ACE2 Ser-680 phosphorylation might crosstalk with S1PR signaling to synergistically inhibit angiogenesis via inhibition of JunB.

In conclusion, this study demonstrates a critical role of PTMs of ACE2, specifically Ser-680 dephosphorylation/Lys-788 ubiquitination in pathogenic angiogenesis. In response to angiogenic cues, AP-1 activation by these PTMs of ACE2 comprehensively upregulates pro-angiogenic genes in ECs. These new findings suggest that targeting Ser-680 dephosphorylation and Lys-788 ubiquitination of ACE2 seems effective in perturbing pathologic angiogenesis involved in diseases such as proliferative retinopathies, tumor angiogenesis, and COVID-19 vasculitis.

## Non-standard Abbreviations and Acronyms

ACE2: angiotensin-converting enzyme 2
Ad-: adenovirus
AMPK: AMP-activated protein kinase
AP-1: activator protein-1
AT1R: Ang II type 1 receptor
ATAC-seq: assay for transposase-accessible chromatin with high-throughput sequencing
ChIP-seq: chromatin immunoprecipitation sequencing
DEG: differentially expressed gene
ECs: endothelial cells
EdU: 5-ethynyl-2-deoxyuridine
EMSA: electrophoretic mobility shift assay
HUVECs: human umbilical vein endothelial cells
MDM2: murine double minute 2
OIR: oxygen-induced retinopathy
PTMs: post-translational modifications
PVP: polyvinylpyrrolidone
RAS: renin-angiotensin system
ROP: retinopathy of prematurity
TFs: transcription factors

## Acknowledgments

We thank Drs. Baochang Lai, Qian Yin, Wenbo Yang, Xiaoli Qian, Peining Liu at Xi’an Jiaotong University and Anne C Lyons and Prof. Jin Zhang at the University of California, San Diego for their technical assistance, discussion, and consultation.

## Sources of Funding

This work was supported by the National Key Research and Development Program of China (2021YFA1301200); the National Natural Science Foundation of China (82430019, 82200455, 92049203); the Key Research and Development Program of Shaanxi (2025SF-YBXM-008); and the Fundamental Research Funds for the Central Universities (xzy022024019).

## Disclosures

None.

## Author Contributions

Y.L., Y.X., X.W., J.Y.-J.S., and Z.-Y.Y. conceived of the study and designed the experiments. Y.L., Y.X., Y.L., Q.L., T.-Y.W.W., B.L., F.H., Z.W., L.W., C.W., W.H., and J.Z. performed the experiments. Y.L., Y.X., and F.H. conducted in silico analysis. Y.L., Y.X., F.H., J.Z., J.Z., Y.X., X.W., J.Y.-J.S., and Z.-Y.Y. analyzed the data and provided the discussion. Y.L., Y.X., X.W., J.Y.-J.S., and Z.-Y.Y. wrote the manuscript. All authors reviewed the paper and approved the final draft.

## Notes

### Competing Interest Statement

The authors have declared no competing interest.

